# Incomplete resilience of a shallow lake to a brownification event

**DOI:** 10.1101/658591

**Authors:** G. Kazanjian, S. Brothers, J. Köhler, S. Hilt

**Affiliations:** Leibniz-Institute of Freshwater Ecology and Inland Fisheries (IGB), Department of Ecosystem Research, Müggelseedamm 301, 12587 Berlin, Germany; Utah State University, Department of Watershed Sciences and Ecology Center, 5210 Old Main Hill, Logan, Utah, 84322-5200, USA

**Keywords:** Dissolved organic carbon (DOC), phosphorus, primary production, ecosystem metabolism, phytoplankton, periphyton, regime shift, water-level change, precipitation, temperate climate

## Abstract

Dissolved organic carbon (DOC) concentrations in many freshwater ecosystems of the northern hemisphere have increased in recent decades due to additional terrestrial inputs. This phenomenon, known as brownification, can strongly alter the physical, chemical, and biological traits of aquatic ecosystems. Extreme rainfall can also cause sudden brownification, known as blackwater events in rivers, while longer term effects on lakes are unknown. Here, we investigated the resilience of a small, temperate, shallow lake to a strong natural flooding-induced brownification event in 2011-2012. From initial DOC and total phosphorus (TP) concentrations of ~12 and 0.04 mg L^−1^, respectively, the lake rapidly reached peak DOC and TP concentrations of 60 and 0.35 mg L^−1^, respectively. By the following year, water levels had returned close to initial values, yet two additional years of monitoring (until summer 2015) and a more recent sample in spring 2019 showed that the lake did not fully return to its pre-brownification state. Instead, DOC and TP concentrations plateaued at concentrations respectively 1.5-fold and twofold greater than pre-brownification values within less than two years and remained at these concentrations in spring 2019. During this initial recovery period the lake exhibited a decline of phytoplankton and a partial recovery of summer periphyton biomass and production, albeit a full return to pre-brownification values was not recorded in either case. DOC and TP concentrations were positively correlated to phytoplankton biomass and negatively to periphyton. As increases in phytoplankton production outpaced decreasing periphyton production, the net result of this brownification event has been an increase in whole-lake areal summertime primary production. This incomplete resilience to a flooding-induced brownification event implies consequences for several ecological and biogeochemical functions of shallow lakes that warrant further investigation and might contribute to the gradual increase of freshwater DOC concentrations in the northern hemisphere.

## Introduction

Dissolved organic carbon (DOC) concentrations in lakes and rivers have increased over the past decades in many regions (Evans et al., 2006; Williamson et al., 2015), mostly due to additional terrestrial inputs (Monteith et al., 2007; Solomon et al., 2015). This has led to brownification becoming a common phenomenon, especially in the northern hemisphere (Roulet and Moore, 2006). Increasing DOC concentrations can significantly impact the chemical, physical, and biological traits of aquatic ecosystems (Brothers et al., 2014; Jones and Lennon, 2015; Solomon et al., 2015; Hedström et al., 2017). Terrestrial organic carbon (OC) inputs contribute to basal resource availability (Pace et al., 2004; Solomon et al., 2011), but they can also reduce primary productivity via shading effects on phytoplankton and periphyton (Karlsson et al., 2009; Thrane et al., 2014). Brownification can also physically alter small lakes by increasing the stability of thermal stratification, with multiple potential effects on algal biomass and productivity (Fee et al., 1996; Houser, 2006; Brothers et al., 2014). Above a concentration threshold of 5-15 mg L^−1^, the negative influence of DOC shading on autochthonous primary production exceeds the positive effects of DOC on resource availability via direct supply of fixed C and potential fertilization of autochthonous production (Karlsson et al. 2009; Jones et al., 2012; Seekell et al., 2015; Kelly et al. 2018). Additionally, increased DOC concentrations can alter phytoplankton species composition and diversity (Lenard and Ejankowski, 2017; Urrutia-Cordero et al., 2017), which in turn can affect the aquatic food web (McGowen et al., 2005).

Apart from a gradual upward trend in DOC concentrations in many freshwaters, DOC inputs and concentrations can fluctuate significantly on shorter timescales. In lowland river systems, sudden “blackwater” events commonly occur when flooding follows prolonged dry periods, transporting high quantities of accumulated terrestrial organic material. Raymond and Saiers (2010) calculated that 86% of the annual DOC flux in small forested catchments occurred in association with rising or falling stream-water hydrographs. The released DOC can lead to severe anoxia in streams and rivers, killing aquatic animals (e.g., Hladyz et al., 2011; Ning et al., 2015). Extensive flooding in the Murray–Darling Basin (Australia) following a decade of drought mobilized several hundred thousand tons of DOC and the plume of hypoxic water affected roughly 2000 km of river channel for up to 6 months (Whitworth et al., 2012). Nonetheless, blackwater events in rivers are often short-lived due to flushing, allowing a rapid recovery of both water quality and the affected fauna (Burford et al., 2008; Kerr et al., 2013).

Flooding-induced brownification events can also occur in lakes, although few cases have been described in the literature. Boreal lakes with a water retention time between one and three years are particularly vulnerable to precipitation-induced browning as climate change models predicting increasing precipitation in this region indicate that many of these lakes will continue to experience browning in the foreseeable future (Weyhenmeyer et al., 2016). As with streams, brownification in lakes can lead to anoxia and have strong effects on water chemistry, algal community composition, biomass and productivity, as well as the mortality of macrozoobenthos and fish (Sadro and Melack, 2012; Brothers et al., 2014; Lenard and Ejankowski, 2017). Due to longer water residence time in lakes, the effects of sudden brownification events on water chemistry and biota are expected to last longer than in rivers. In lakes, DOC removal is primarily facilitated by microbial mineralization (Hanson et al., 2011), flocculation (von Wachenfeldt and Tranvik, 2008), and photolytic mineralization (Granéli et al., 1996; Cory et al., 2014). The resilience of these systems, here defined as the rate of recovery after a disturbance (Tsai et al., 2011), to sudden brownification events (short-term increases in DOC loading) remains unknown.

In this study, we analyzed the resilience of a small, temperate, shallow lake to a sudden natural brownification event previously described by Brothers et al. (2014). Due to high precipitation and naturally rising water levels, the DOC concentrations in this lake had increased five-fold, from ~12 mg L^−1^ in 2010 to a maximum of about 60 mg L^−1^ by 2012.Concurrently with increasing DOC concentrations, total phosphorus (TP) and iron (Fe) concentrations had risen dramatically. Benthic and planktonic primary producers had exhibited opposing responses, with an increase in phytoplankton gross primary production (GPP) due to increased nutrient availability (due largely to internal loading, likely linked to exacerbated thermal stratification), while periphyton biomass strongly declined, likely due to shading by phytoplankton, DOC, and Fe and substantially cooler hypolimnetic temperatures (Brothers et al., 2014). We continued the examination of lake water quality and primary production for three years following peak DOC concentrations (until summer 2015), to investigate their potential recovery. We also investigated the response of phytoplankton community composition to the initial increase in DOC concentrations and over the subsequent years. Along with declining water levels after 2012, we anticipated a reduction of external and internal DOC loading, producing a gradual decrease in lake DOC and TP concentrations. Accordingly, we hypothesized that pelagic GPP and phytoplankton community composition would return to pre-brownification conditions, driven by decreasing nutrient concentrations and deeper (pre-brownification) mixing depths, while benthic primary production would recover due to increased light availability.

## Methods

### Study site

Kleiner Gollinsee (hereafter referred to as Gollinsee) is a small (0.03 km^2^), shallow (Z_mean_: 1.7 m, Z_max_: 2.9 m; 2010 values), eutrophic lake located in northeastern Germany (53°01’N, 13°35’E). The lake lacks surface in- or outflows. A net gain of 133 mm of groundwater was previously recorded from June 8 to October 19, 2010 (Rudnick, 2011) indicating a water residence time of approximately 5 years. Gollinsee is sheltered from strong winds by a reed belt (*Phragmites australis* Trin. ex Steud.) and surrounding alder trees (*Alnus glutinosa L.*) and lacks submerged macrophytes (Brothers et al., 2013).

From November 2010 to November 2014, as part of an unrelated experiment tracing terrestrial particulate organic carbon within the aquatic food web (Attermeyer et al., 2013; Scharnweber et al., 2013), Gollinsee was divided into two similarly-sized basins using a plastic curtain. The data shown in this study represent whole lake averages for the years 2010 and 2015 and separate averages for each basin, hereafter referred to as north and south basins, during the years the lake was split (2011 – 2014). Water quality parameters in 2010 reflect the lake’s pre-brownification (baseline) state (further supported by measurements taken in 2007), 2011 marks the onset of the flooding and brownification event which reached its peak in summer 2012 (Brothers et al., 2014), and 2013 is considered to represent the beginning of the lake’s recovery period from the flooding and brownification event.

### Water sampling and water quality analysis

Integrated water samples (every 0.5 m from the water surface to just above the sediment) were retrieved using a Ruttner-like water sampler about every three months from spring 2013 to summer 2015. A single, final water sample was taken on April 27^th^, 2019. Water samples were transported to the laboratory in cooling boxes and water chemistry parameters (listed below) were analyzed the following day. During stratified periods, separate integrated samples were collected from the epilimnion and hypolimnion. Mixing depths were determined using vertical profiles of dissolved oxygen (O_2_) concentrations, pH, and water temperature, measured by a Yellow Springs Instruments (YSI) multi-probe sonde. In addition, a lake-center weather station recorded and transmitted global radiation, wind speed, and air temperature data every 30 minutes over the same period. Due to technical problems with the weather station, data from January to mid-September 2013 and all of 2015 were retrieved from a nearby weather station (at Döllnsee, 3.5 km southeast of Gollinsee). Water column light attenuation was measured every field campaign using two Underwater Spherical Quantum Sensors (LI-193, LI-COR) deployed 50 cm apart. Water level was measured monthly by local conservation authorities (kindly provided by R. Michels, Biosphärenreservat Schorfheide-Chorin).

We analyzed water samples for concentrations of total phosphorus (TP), total dissolved phosphorus (TDP), soluble reactive phosphorus (SRP), and dissolved nitrogen (DN) following German standard procedures (DEV, 2009). We calculated particulate phosphorus (PP) by subtracting the values of TDP from TP. Dissolved organic phosphorus (DOP) was calculated by subtracting SRP from TDP values. DOC concentrations were measured with a total organic carbon (TOC) Carbon-Analyzer (TOC 5000, Shimadzu), and iron (Fe) concentrations were analyzed using an inductively-coupled plasma optical emission spectrometer (ICP-OES) with an inductively-coupled argon plasma unit (iCAP 6000-Duo, Thermo Fisher Scientific). Furthermore, to explore the effects of DOC and humic substances on light attenuation in the water column, we compared the fluorescence of filtered lake water at 470 nm, measured using a pulse amplitude modulated fluorometer (Phyto-PAM, Walz, Effeltrich, Germany), with lake water DOC concentrations.

### Biomass and production of phytoplankton and periphyton

For phytoplankton biomass, aliquots of the water samples were filtered onto six 25 mm Whatman Glass Fiber Filters (GF/F), three of which were used to determine dry weight and C:N ratios, while the rest were used to measure chlorophyll *a* (chl-*a*) concentrations via high performance liquid chromatography (HPLC) following Shatwell et al. (2012). Rapid photosynthesis-irradiance (P-I) curves of phytoplankton were measured using a Phyto-PAM fluorometer following a dark adaptation period of at least 15 minutes. Chl-*a* concentrations were paired with P-I parameters, global radiation, and water column light attenuation to calculate phytoplankton GPP following Brothers et al. (2013). Furthermore, using calibration files based on specific phytoplankton groups sampled from Müggelsee (Berlin, Germany), we used fluorescence measurements at four wavelengths to identify the percent contribution of diatoms, green algae, and cyanobacteria to phytoplankton biomass(data from July 28^th^, 2011 excluded due to equipment malfunction).

From 2010 to 2014, we incubated artificial plastic substrates in Gollinsee for a month between June and July to measure summer periphyton biomass accumulation and GPP. During this period, GPP approaches maximum values in shallow lakes (Liboriussen and Jeppessen, 2003; Brothers et al., 2013). Upon harvest, the plastic substrates were transported to the lab in a dark, humid cooler, where rapid photosynthesis-light curves of periphyton were measured using a Phyto-PAM Emitter Detector Fiberoptics (EDF) unit. Periphyton was subsequently scraped from the artificial substrates using a toothbrush and filtered lake water, and filtered onto 25 mm Whatman GF filters to determine chl-*a* content and C:N ratios, following the protocol described above for phytoplankton.Chl-*a*, P-I parameters, global radiation, vertical light attenuation, and the depth distribution of the lake were used to calculate periphyton GPP (see Brothers et al. (2013) for details).

### Statistical analyses

Spearman’s rho correlation indices were calculated to test for a relationship between background lake water fluorescence and DOC concentrations and between primary producer biomass or production and DOC or TP concentrations in the water column. All statistical analyses were performed using R version 3.4.2 (R group).

## Results

### Lake water parameters and quality

After a strong increase in the water level of Gollinsee between 2011 and 2012, a gradual decline in water levels following summer 2013 returned the water surface to roughly pre-flood levels by the summer of 2015 (~ 20cm higher than mean water levels from 2007-2010; Fig. S1). Lake water DOC concentrations, having also reached maximum values during the summer of 2012, decreased more rapidly than water levels (Fig. 1A). Within one year following peak values (i.e. by summer 2013), DOC concentrations had already fallen by roughly 40%, even though water levels had decreased only marginally (5%) over that same period. Thereafter, the decline in DOC concentrations slowed down (declining another 29% by summer 2014) despite a stronger concurrent water level decline (19%), while in the last year of sampling, the DOC concentration remained at roughly 17.5 mg L^−1^, ~50% greater than 2010 pre-brownification values.

**Fig. 1:**
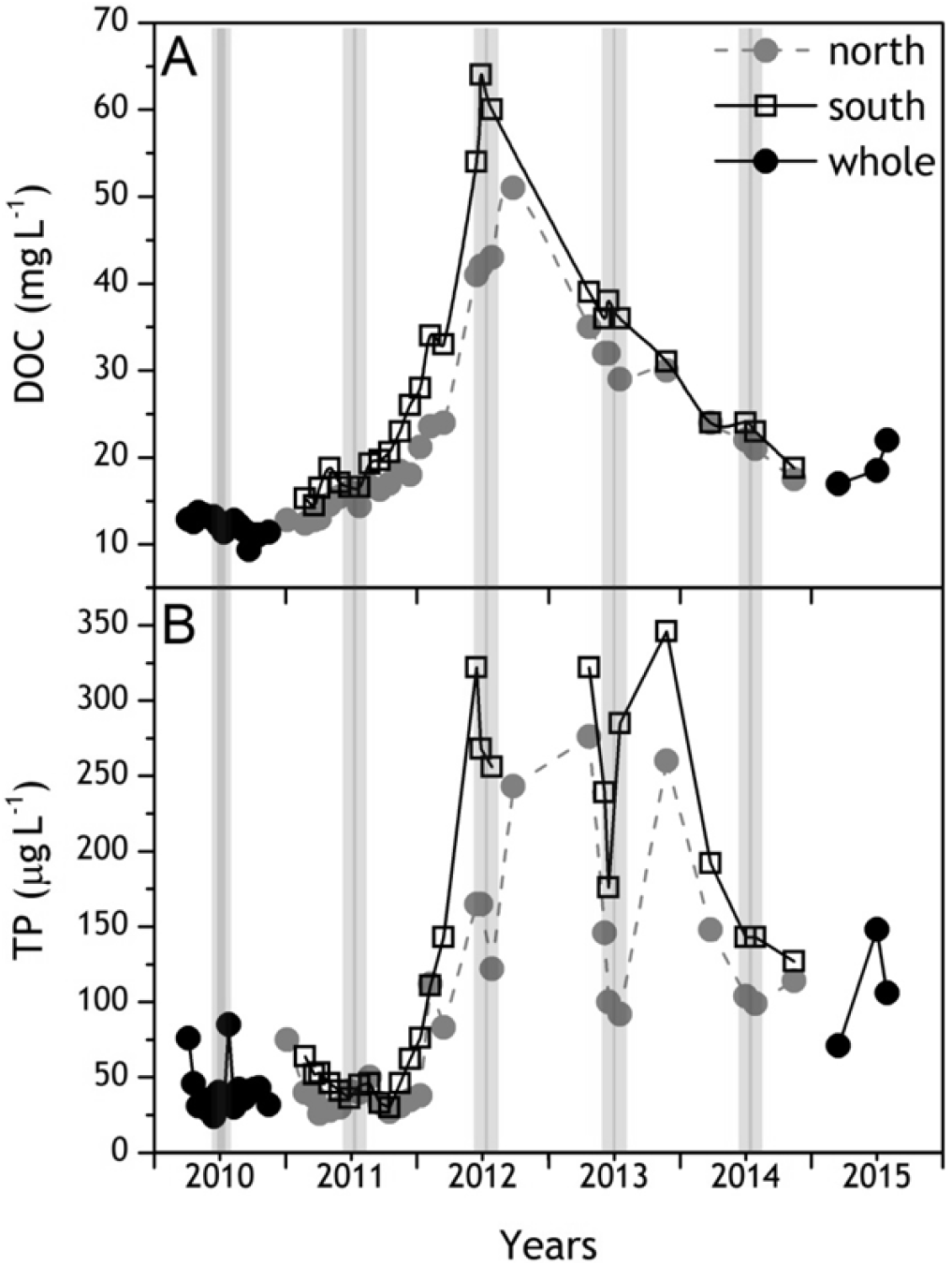
Dissolved organic carbon (DOC, mg L^−1^) and total phosphorus (TP, μg L^−1^) concentrations in Lake Gollinsee, from 2010 to 2015. 2010 and 2015 values represent that of the whole lake, whereas 2011-2014 values are shown for the two sides when the lake was split in half. Bars indicate annual benthic production sampling times (June-July). Data from 2010-2012 are taken from Brothers et al. (2014).

Following an initial strong increase in 2011, concentrations of TP fluctuated greatly through 2012 until the end of 2013 but were generally lower thereafter (Fig. 1B) and seemed to have stabilized by the end of 2014 at concentrations around double those measured prior to brownification (127 μg L^−1^ in July 2015, compared to 58.5 μg L^−1^ in 2010). An additional field excursion and water sample taken in April 2019 confirmed that the water level (46 cm, Fig. S1) as well as DOC (21.6 mg L^−1^) and TP (131 μg L^−1^) concentrations have not declined since 2015. Concentrations of SRP (Fig. 2A), which had also risen dramatically in summer 2012, exhibited their highest peak (261 μg L^−1^) in the southern basin in autumn 2013. This peak was preceded by peaks in PP (Fig. 2B), and in DOP concentrations (Fig. 2C) in summer 2013, albeit peak DOP concentrations coincided with peak DOC concentrations (summer 2012). Concentrations of DN and Fe (Fig 2D, F) also exhibited a similar trend to DOC, while trends in ammonium concentrations followed those of SRP concentrations (Fig. 2E). Concentrations of Mn increased significantly with maxima recorded in summer 2012 (Fig. 2G). Furthermore, all of SRP, PP, DN, NH_4_, Fe, and Mn concentrations exhibited peaks at least in one of the basins in autumn 2013 (Fig. 2). In general, the two lake basins showed comparable dynamics for all parameters with only slight differences in peak concentrations. To establish a longer-term perspective of the water quality in Gollinsee prior to the brownification event, we also report data from a prior sampling campaign conducted in 2007 (Supplementary Table1).

**Fig. 2:**
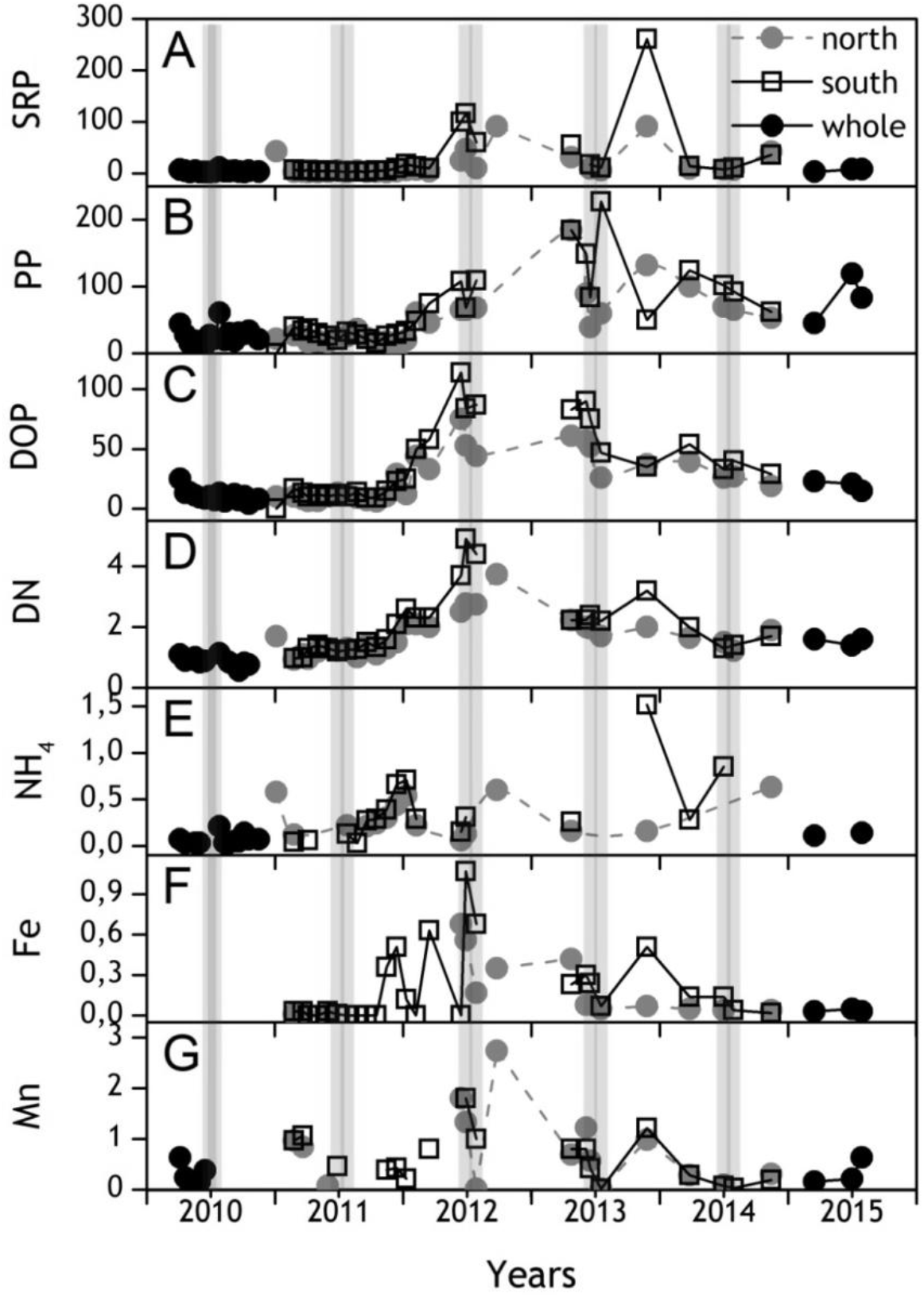
Concentrations of (A) soluble reactive phosphorus (SRP, in μg L^−1^), (B) total particulate phosphorus (PP = TP – TDP, in μg L^−1^), dissolved organic phosphorus (DOP = TDP -SRP, in μg L^−1^), dissolved nitrogen (DN, in mg L^−1^), ammonium (NH_4_^+^, in mg L^−1^), iron (Fe, in mg L^−1^), and manganese (Mn, in mg L^−1^) in Lake Gollinsee from 2010 to 2015. 2010 and 2015 values represent that of the whole lake, whereas 2011-2014 values are shown for the two sides when the lake was split in half. Grey bars indicate annual benthic production sampling times.

### Light availability

Mean global radiation values measured at the surface of Gollinsee during our periphyton study periods (June-July) dropped in 2011 but increased gradually every year from 2011 to 2014 (Fig. 3A). Light attenuation values were highest in 2012 (Fig. 3B), leading to the most reduced euphotic zone depth during that same year (Fig. 3C). Thereafter, despite increasing light conditions every year, the water column did not fully return to its pre-brownification light attenuation levels (1.8 m^−1^ in 2010 *vs* 2.9 m^−1^ in 2015) and euphotic zone depth (2.6 m in 2010 *vs* 1.6 m in 2015) (Figs. 3B, C). The background fluorescence of filtered water (i.e. likely caused by colored humic substances) was higher in 2012 and 2013 than in previous years, but values in 2014 were similar to those in 2011 (Fig. S2), and values had returned to pre-brownification levels by 2015. Background fluorescence and DOC concentrations were strongly correlated (Spearman’s rho = 0.958, *P* = 0.0002, Fig. S2).

**Fig. 3:**
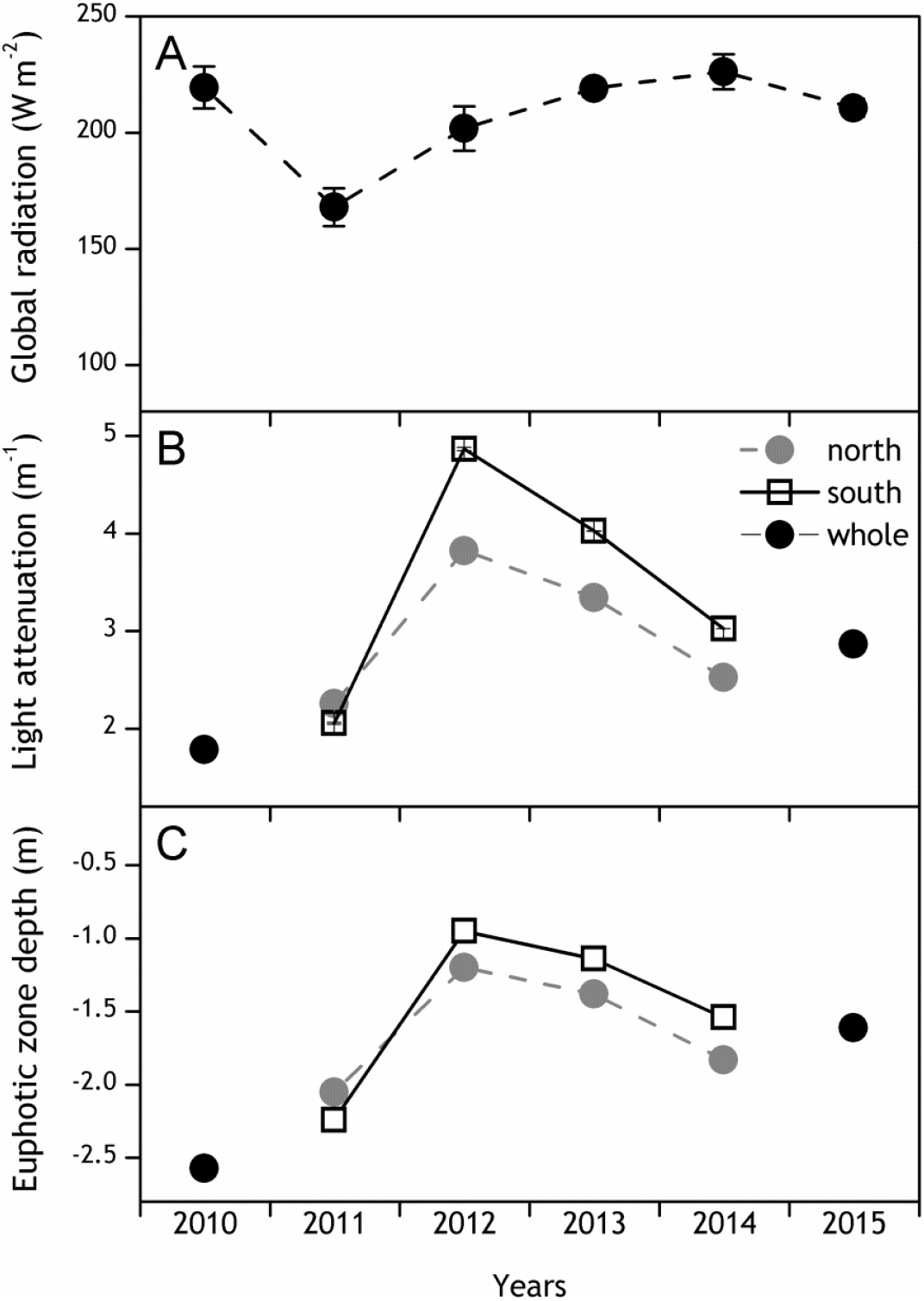
Differences in global radiation at the water surface (A, ± standard error), water column light attenuation levels (B, ± standard error) and euphotic zone depth, defined as 1% of PAR (C) at Lake Gollinsee in June and July between 2010 and 2015. Global radiation was measured continuously at regular intervals of 10 minutes in 2011, hourly in 2012, and every 30 minutes in 2014. On-site 2013 and 2015 data are lacking due to weather station malfunction, and we thus show global radiation measurements from Döllnsee instead (3.5 km from Gollinsee). Light attenuation values represent the average of two direct measurements per year, at the start and end of the studied period.

### Biomass and production of phytoplankton and periphyton

Throughout the study, Gollinsee was dominated by phytoplankton production, though the biomass and GPP of the primary producers varied between years and with changes in DOC and TP concentrations (Figs. 4, 5). Phytoplankton biomass reached its peak in 2013 (191 mg chl-*a* L^−1^ in the epilimnion of the southern basin), before decreasing to pre-brownification (2010) levels by 2014.Phytoplankton GPP rates peaked at 4.9 g C m^−2^ d^−1^ in the southern basin in 2013 and dropped three-fold the following year. Phytoplankton biomass was positively correlated with TP. In contrast, periphyton biomass and GPP showed an inverse relationship to DOC and TP concentrations (Table 1; Fig. 5) and thus were at their lowest during peak brownification. Regarding phytoplankton community composition (Fig. S3), diatoms dominated the phytoplankton community before the brownification event. During the first summer after the onset of brownification (2011), green algae represented more than three-quarters of the phytoplankton biomass in the two basins of the lake. In the two subsequent years, diatoms established the majority of phytoplankton biomass, followed by a more heterogeneous composition in 2014. During the summer of 2015, after the removal of the curtain splitting the lake, the phytoplankton community was dominated by cyanobacteria.

**Fig. 4:**
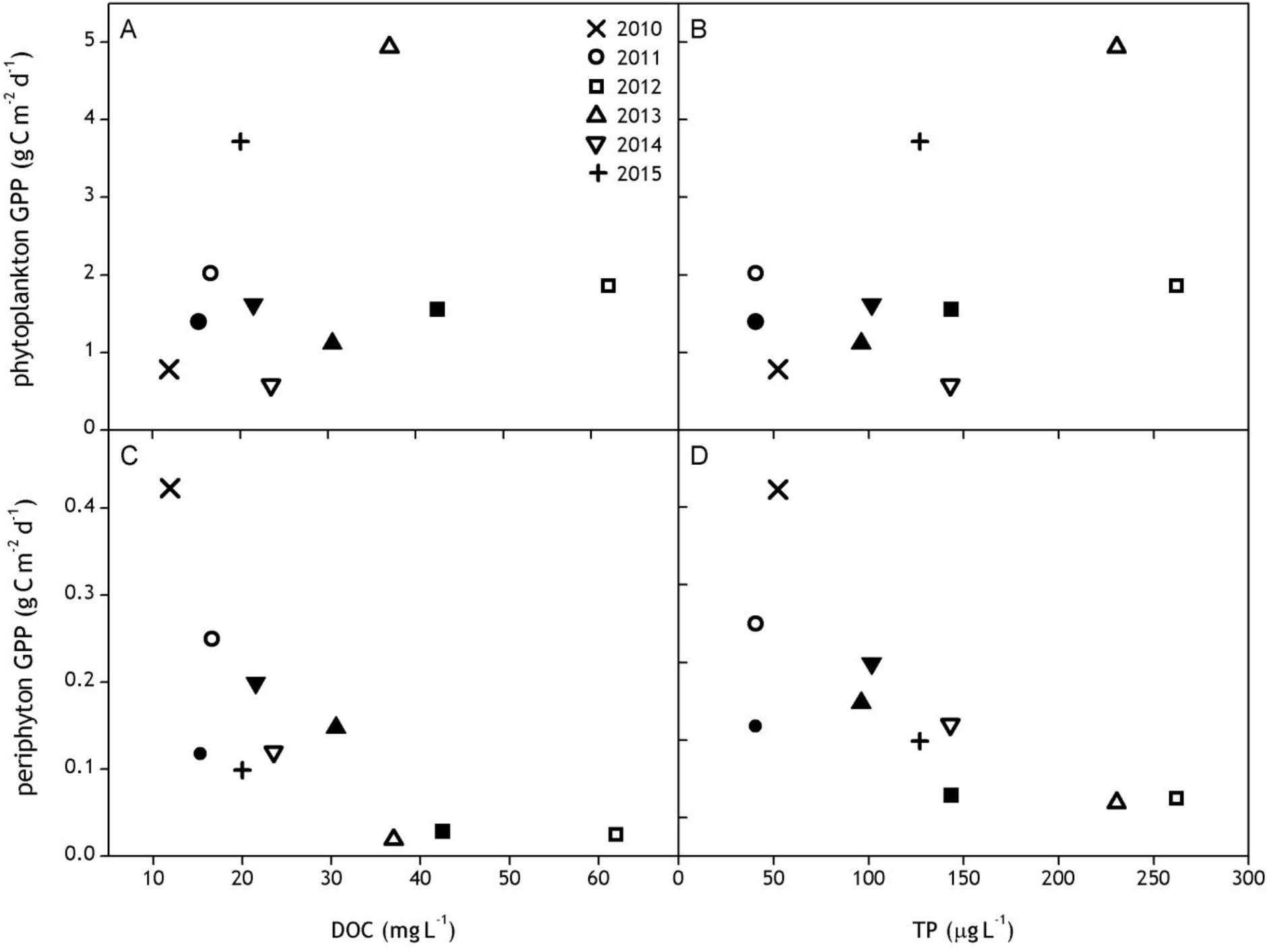
Summer GPP (phytoplankton in top row and periphyton GPP in bottom; in g C m^−2^ d^−1^) and water DOC (in mg L^−1^) and TP (in μgL^−1^) concentrations in Lake Gollinsee from 2010 to 2015. Filled symbols represent values from the northern basin, empty symbols correspond to values from the southern basin. A single symbol from each of 2010 and 2015 represents whole-lake calculations.

**Fig. 5:**
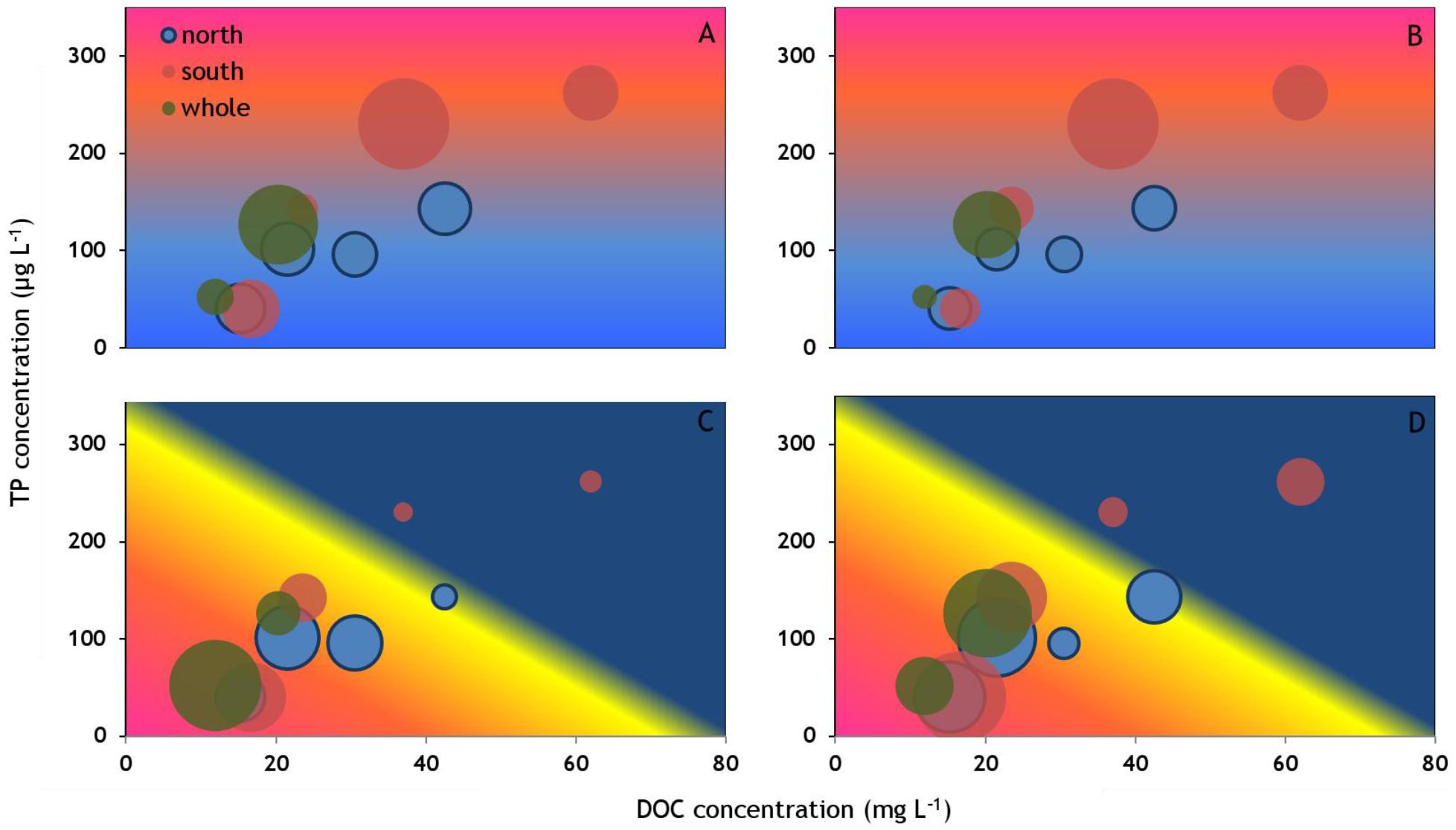
Hypothesized (background) and measured (circles) summer phytoplankton gross primary production (GPP) and biomass (A and B, respectively) and epilimnetic periphyton GPP and biomass (C, D) in Lake Gollinsee, in relation to water column DOC and TP concentrations from 2010 to 2015. The colored background (concept taken from the model of Vasconcelos et al, 2016) indicates a gradual (A, B) or sudden (C, D) shift from favorable (red) to limited (blue) growth conditions. The size of the circles corresponds to the relative value of GPP or biomass. Blue circles represent measurements from the northern basin, red circles from the southern basin, and green circles from the whole lake during the years when the curtain splitting the lake was not deployed.

**Table 1:** Spearman’s correlation indices and *P*-values of phytoplankton and periphyton biomass and GPP values with water dissolved organic carbon (DOC) and total phosphorus (TP) concentrations in Gollinsee between 2010 and 2015. Significant values are represented in bold.

|  | DOC |  | TP |  |
| --- | --- | --- | --- | --- |
|  | Spearman's rho | *P*-value | Spearman's rho | *P*-value |
| Phytoplankton biomass | 0.527 | 0.123 | 0.748 | **0.013** |
| Phytoplankton GPP | 0.236 | 0.514 | 0.300 | 0.403 |
| Periphyton biomass | -0.745 | **0.018** | -0.651 | **0.042** |
| Periphyton GPP | -0.721 | **0.024** | -0.784 | **0.007** |

## Discussion

The shallow, temperate lake we studied for five years did not exhibit a full recovery from a brief (one-year) but strong, flood-induced brownification event. Three years after reaching peak levels (60 mg L^−1^), DOC concentrations were still 1.5-fold greater than pre-brownification values. The decrease of TP concentrations was even less pronounced, remaining at more than double the baseline concentrations by the end of this study. These concentrations seem to have plateaued thereafter, as indicated by comparable DOC and TP concentrations measured nearly four years after the conclusion of this study (spring 2019). Summer phytoplankton biomass and GPP thus remained elevated, while periphyton biomass and GPP, being negatively correlated with both DOC and TP concentrations, remained lower than pre-brownification levels.

### DOC and TP dynamics

Along with the water-level rise at Gollinsee (Fig. S1), the sharp five-fold increase in DOC concentrations (from early 2011 to the summer of 2012) was attributed to DOC leaching from the surrounding flooded degraded peatlands (through most of 2011 and early 2012), followed by internal loading due to the reductive dissolution of iron-bound DOC in the lake sediments by mid-2012 (Brothers et al., 2014). The subsequent decrease in DOC concentrations, roughly by 30 mg L^−1^ a^−1^ in the first year after peak DOC concentrations (2012-13), was faster than the gradual water-level drop, indicating that water levels and DOC dynamics were at least partially decoupled in this lake. The decline in DOC concentrations was likely due to active removal of the DOC from the water column as well as diminishing DOC loading rates into the water column. Active removal mechanisms include bacterial (von Wachenfeldt and Tranvik, 2008) and photolytic mineralization (Granéli et al., 1996; Bertilsson and Tranvik, 2000), as well as flocculation resulting in burial in the sediments (von Wachenfeldt and Tranvik, 2008; Skoog and Arias-Esquivel, 2009). Concerning the specific rates of DOC removal possible by different mechanisms, rising DOC concentrations can increase bacterial DOC consumption rates by up to 68% when nutrients are not limiting (measured in nearby Lake Schulzensee; Attermeyer et al., 2014), with consumption rates reaching 32. 1 mg C L^−1^ a^−1^ in eutrophic lakes featuring elevated bacterial growth efficiencies (Biddanda et al., 2001). Photolytic mineralization is also known to be an effective pathway for removing terrigenous organic carbon (Obernoster and Benner, 2004), with estimations (based on Bertilsson and Tranvik, 2000) suggesting that it could explain about half of our observed decrease in DOC concentrations (9.4 mg C L^−1^ a^−1^, using the mean global radiation at Gollinsee for the year 2012 and assuming the top 2 cm water layer to have been subject to photolysis). Other processes such as flocculation, might also contribute to DOC removal from the water column.

Regarding diminishing DOC loading rates into the water column, both external and internal DOC loading likely diminished following 2012. External (terrestrial) DOC leaching from the surrounding landscape would decline naturally with returns to lower water levels. Concomitant decreases of DOC and Fe concentrations (Figs. 1A, 2F) indicate that the co-precipitation of DOC with iron-containing minerals (namely iron sulfide) could have also played a major role in DOC removal(Skoog and Arias-Esquivel, 2009), and a return to greater DOC storage in the lake sediments. Prevailing anoxic conditions during peak brownification (summer 2012) would have presumably driven sulfide concentrations to increase in the water column, which in turn would have led to iron sulfide precipitation. Although we did not directly measure sulfide concentrations in the water column, the sediment surface in Gollinsee post-brownification was characterized by a fluffy black precipitate (pers. obs.), a known characteristic of sulfide. An increasing proportion of oxic sediment layers during benthic periphyton recovery (see below) might have produced a positive feedback loop inhibiting the re-release of DOC from the sediments (Peter et al., 2017), while also intercepting the release of nutrients from the sediment (Vasconcelos et al., 2016). After 2013, DOC concentrations declined more slowly and eventually stabilized at concentrations which were roughly 50% greater than pre-brownification values.

Total phosphorus concentrations exhibited a longer delay in recovery than DOC concentrations (Fig. 2). Phosphorus released from both catchment soils and sediments during anoxic conditions was likely taken up by phytoplankton when pelagic production was boosted during the brownification event. This is corroborated by the highest recorded values of PP (in 2013 in the southern basin, Fig. 2) coinciding with the highest phytoplankton GPP (Fig. 4). The assimilation of P in phytoplankton potentially ensured that it did not co-precipitate with iron when the water column became oxic again, as indicated by a lack of correlation between TP and Fe concentrations during the recovery phase. Other recorded peaks during autumn 2013 in concentrations of SRP, Fe, and Mn (Fig. 2) were likely caused by the mixing of the nutrient-rich and anoxic hypolimnion with the epilimnion following a strong summer stratification (Fig. S4).

### Response of primary producers to brownification

Phytoplankton and periphyton biomass and GPP responded in opposing manners to the brownification event and through the lake’s recovery. Phytoplankton GPP was enhanced by brownification due to higher TP concentrations and compressed mixing depths, exacerbating the shading of periphyton by DOC and Fe (Jones, 1992; Brothers et al., 2014). Previous studies have reported a similar rise in pelagic GPP during brownification events, likely due to an increase in phosphorus availability (Grabowska et al., 2003; Zwart et al., 2016). Browning also alleviates pelagic algal nutrient limitation by shading benthic competitors and preventing them from intercepting the release of nutrients from the sediments (Vasconcelos et al., 2016). Consequently, light extinction (which limits GPP) and nutrient availability (which stimulates GPP) are non-linearly related to DOC concentration (Seekell et al., 2015; Kelly et al. 2018). This is also demonstrated in our results (Fig. 5), which strongly support previous theoretical model predictions on the differential response of pelagic and benthic primary producers to increasing DOC and TP concentrations (Vasconcelos et al., 2016). Higher DOC and TP concentrations coincides with a gradual increase in phytoplankton biomass and production (Fig. 5A, B), as well as an increasing light attenuation within the water column that diminishes benthic GPP. This trend continues until crossing a threshold (yellow background line in Fig. 5C, D) beyond which benthic algae are no longer productive. Since light attenuation is driven by both DOC and phytoplankton biomass, which is itself correlated to TP concentrations within the water column (Table 1), this threshold varies along a DOC : TP concentration spectrum. In contrast, if water quality parameters return to pre-brownification levels, the lower DOC and TP concentrations enhance and limit periphyton and phytoplankton GPP, respectively.

While our study confirms published theoretical models, it also covers a much wider spectrum of DOC concentrations than previously studied. Most studies to date have focused on DOC concentrations up to 20 mg L^−1^. In our study, due to the occurrence of the strong natural brownification event, DOC concentrations increased to almost three-fold those amounts, providing a beneficial opportunity to extend these theories with additional empirical data. It has been suggested that whole-lake GPP should fall to negligible levels when DOC concentrations exceed a threshold of 15 mg L^−1^ due to extreme shading effects (e.g. Hanson et al 2003; Seekell et al., 2015; Kelly et al. 2018). This study found that high pelagic GPP production persisted well above that threshold (Figs. 4 & 5) due to very shallow mixing depth and low color/DOC ratios. The compression of mixing depths by rising DOC may keep the ratio between euphotic and mixing depth at a level sufficient for intense pelagic GPP only in small, sheltered lakes such as Gollinsee. Light extinction by high DOC concentrations is the underlying reason for limiting GPP (Karlsson et al. 2009), yet DOC can highly differ in its color and light-absorption properties (Pace and Cole, 2002). Fittingly, the response of light attenuation to increasing DOC concentrations observed in this study was substantially lower than has been reported in the literature (Kelly et al. 2018; Fig. S5).

By the end of our study, the lake had experienced only a partial recovery in most measured water quality parameters. Consequently, periphyton GPP never returned to pre-brownification (2010) rates (Fig. 4). Despite the reprieve from DOC and Fe shading following 2012, phytoplankton biomasses continued to fluctuate in the years following the brownification event (Fig. 4), which would continue to suppress benthic algal production, potentially reflecting the establishment of a classic shallow lake stable state relationship between planktonic and benthic primary producers in this lake (e.g., Genkai-Kato et al., 2012). Regarding this sustained phytoplankton production, we noted an initial decline in phytoplankton GPP in 2014, and considered that a subsequent increase in 2015 may have been associated with sediment disturbance due to the removal of the lake division barrier in November 2014. However, water column TP concentrations in 2015 were not significantly greater than those at the end of 2014 (prior to the curtain removal), indicating that the increase in 2015 may instead have been a continuation of the previous unstable phytoplankton dynamics.

Along with changes in production and biomass, we observed significant shifts in phytoplankton composition in our study (Fig. S3), following measured changes in nutrient availability (Berggren et al. 2015; Creed et al. 2018). Prior to brownification, this lake’s community was dominated by diatoms. We had anticipated that increased stratification and internal nutrient loading in 2012 (the peak year of this brownification event) would favor cyanobacterial dominance (Reynolds et al., 1987), yet diatoms instead continued to dominate in 2012, following a brief period of green algal dominance in 2011, corresponding to the initial disturbance period (Fig. S3). However, further long-term observation indicated a gradual relative decline in diatom dominance and by 2015 cyanobacteria had come to dominate the lake’s planktonic primary producer community, even though by this time water levels and water quality were closer to the initial 2010 conditions than in the intervening years. These results indicate that community composition shifts in response to sudden brownification events may only develop gradually, following years after an initial disturbance, and underlining the importance of long-term studies in tracking the effects of disturbances on ecosystems.

It is unclear whether Gollinsee, given more time, will fully return to its pre-brownification state, or whether it has entered a new stable state as a result of the 2012 disturbance. By the end of this study, the proportion of phytoplankton from the overall lake production GPP remained elevated, supported by DOC and TP concentrations that plateaued higher compared to the years prior to brownification. It has been hypothesized that resource pulses can trigger regime shifts with lasting effects on food webs (Holt, 2008), as has apparently occurred in our study lake. Gollinsee was already a phytoplankton-dominated lake in 2010 (Brothers et al., 2013), but the overall increase in phytoplankton GPP during brownification and its incomplete recovery resulted in a higher whole-lake GPP in 2015 compared to 2010. As modelled by Genkai-Kato et al. (2012), the loss in benthic GPP was outpaced by the concurrent increase in pelagic GPP. Oligotrophic, clear lakes have been shown to exhibit a decline in whole-system GPP following an increase in water DOC concentrations (Karlsson et al. 2009; Ask et al. 2009). In such lakes, a minor increase in pelagic GPP may be outpaced by declines in benthic GPP due to DOC and phytoplankton shading. In contrast, eutrophic lakes such as Gollinsee, lacking submerged macrophytes and already dominated by phytoplankton, appear to exhibit a significant increase in pelagic production generated by brownification-triggered internal nutrient loading and shallower mixing depth (Jones, 1992; Brothers et al., 2014). This can compensate for the decrease or complete disappearance of an already-low benthic algal production, driving an overall increase in whole-lake GPP (Genkai-Kato et al., 2012).

The incomplete return of DOC concentrations to pre-brownification levels might also imply that the long-term effects of extreme rainfall events contribute to the general trend of increasing DOC concentrations in freshwater systems of the northern hemisphere. A study of 120 Swedish lakes predicted that an increase in precipitation would result in greater terrestrially-derived DOM concentrations and diminish the influence of in-lake processing on DOM quality (Kellerman et al., 2014). DOC concentrations and its quality can also have significant positive effects on bacterioplankton communities (Crump et al., 2003; Kritzberg et al., 2006) which in turn impact DOC mineralization rates in the system (Attermeyer et al., 2014). With higher frequency of extreme rain events expected in the region (Meehl et al., 2000; van den Besselaar et al., 2012), the trends and impacts of such brownification events will further intensify (de Wit et al., 2016), increasing carbon export from terrestrial to aquatic sources, altering aquatic primary production and greenhouse gas emissions. It is estimated that an increase of 10% in precipitation could lead to a 30% mobilization of OC from soils to freshwaters (de Wit et al., 2016). The increase in DOC concentration in lakes across recent decades has also led to increases in OC burial rates (Anderson et al., 2014), though such impacts caused by short-term brownification events have yet to be thoroughly studied.

We conclude that shallow, eutrophic lakes can exhibit an incomplete resilience to brownification events, as indicated by a lack of full recovery of DOC and nutrient concentrations, as well as planktonic and benthic primary production to pre-brownification levels. With a projected increase in the number of extreme rainfall events coupled to global change, sudden brownification events might become an increasingly common phenomenon, particularly in small shallow lakes. Our findings that short-term weather events can have long-term, sustained effects on lakes and their food webs may thus have far-reaching consequences for trends in water quality and aquatic biogeochemical cycling in the face of ongoing climate change.

## Acknowledgements

We thank S. Meyer, T. Hintze, R. Hölzel, B. Stein, A. Lüder, H. J. Exner, T. Rossoll, and E. Zwirnmann for their technical assistance. We further acknowledge the productive discussions with J. Gelbrecht, T. Goldhammer and M. Kaupenjohann during the writing of this manuscript. R. Michels, H. Mauersberger (Biosphärenreservat Schorfheide-Chorin), R. Mauersberger (Förderverein Feldberg-Uckermärkische Seen e.V.), and R. Tischbier (Stiftung Pro Artenvielfalt) kindly provided background information and lake access.This study was financed by the TERRALAC (http://terralac.igb-berlin.de) and the LandScales project of the Wissenschaftsgemeinschaft Leibniz (WGL).

