## Supplementary for "Incomplete resilience of a shallow lake to a brownification event"

**Table S1**: Mean (+SD) concentrations of dissolved organic carbon (DOC, mg $L^{-1}$), total phosphorus (TP, µg $L^{-1}$), soluble reactive phosphorus (SRP, in µg $L^{-1}$), total particulate phosphorus (PP = TP – TDP, in µg $L^{-1}$), total nitrogen (TN, in mg $L^{-1}$), and ammonium ($NH_4^+$, in mg $L^{-1}$) measured monthly in Gollinsee from March to October 2007 (n = 8). Data taken from the Brennecke (2008).

| DOC | TP | SRP | PP | TN | $NH_4^+$-N |
|---|---|---|---|---|---|
| **12.91 ± 0.56** | 53.13 ± 16.31 | 4.25 ± 1.89 | 38.25 ± 12.9 | 1.42 ± 0.31 | 0.15 ± 0.16 |

**Fig. S1:** Water level fluctuations at the measuring gauge level in Lake Gollinsee between 2007 and 2019

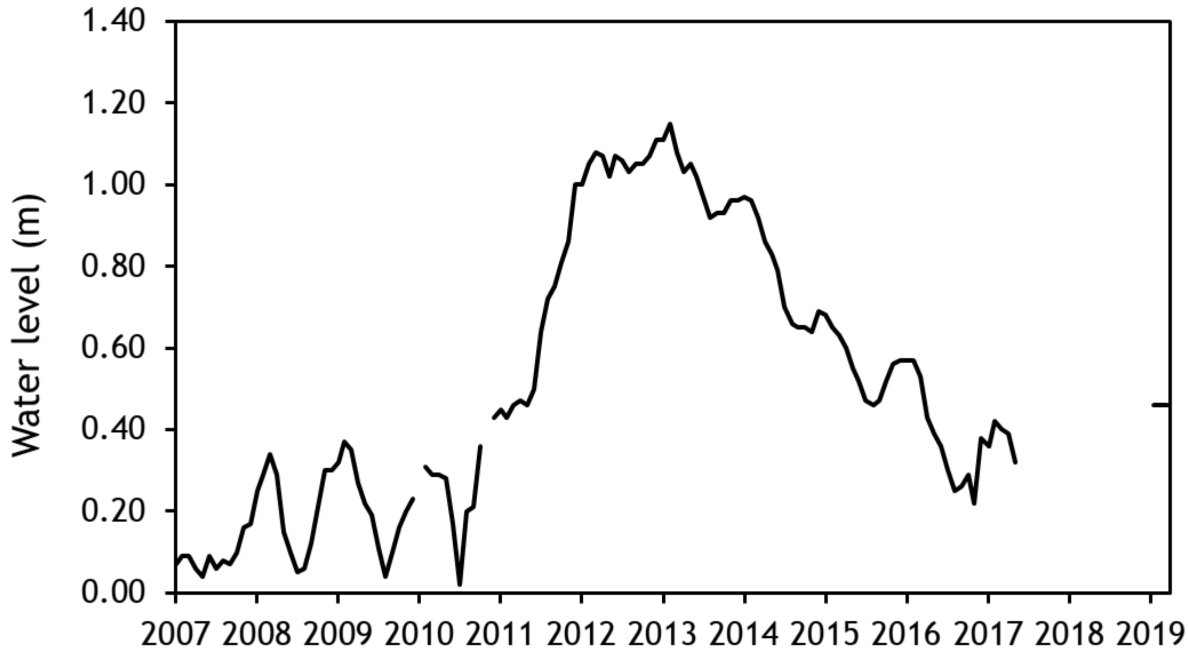

**Fig. S2:** Correlation between DOC concentrations and background water column fluorescence at 470 nm in the whole lake (2010 and 2015) and in two lake sides of split Lake Gollinsee from June and July of 2011 to 2014.

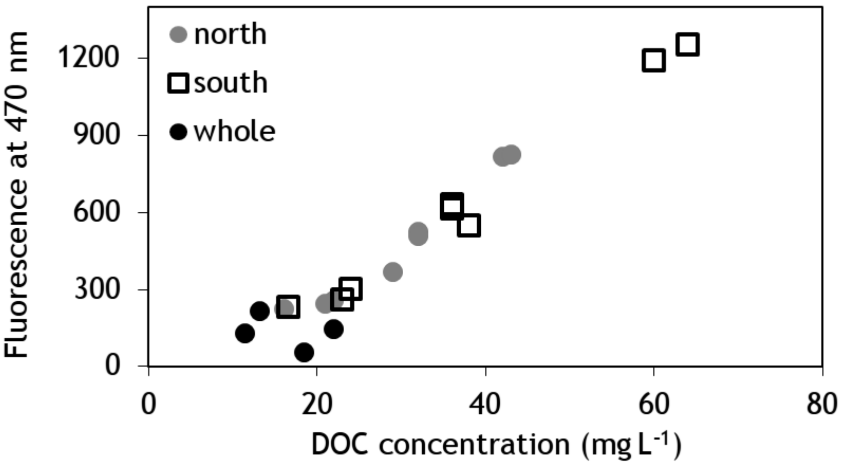


**Fig. S3:** Percentage contribution (average summer values) of the different phytoplankton groups (cyanobacteria, green algae, and diatoms) to total phytoplankton chl-*a* (measured by a PhytoPAM fluorometer) in the northern basin (A), in the southern basin (B), and in the whole lake (C) Gollinsee in summer (June-July) 2010-15 (n= 3-8).

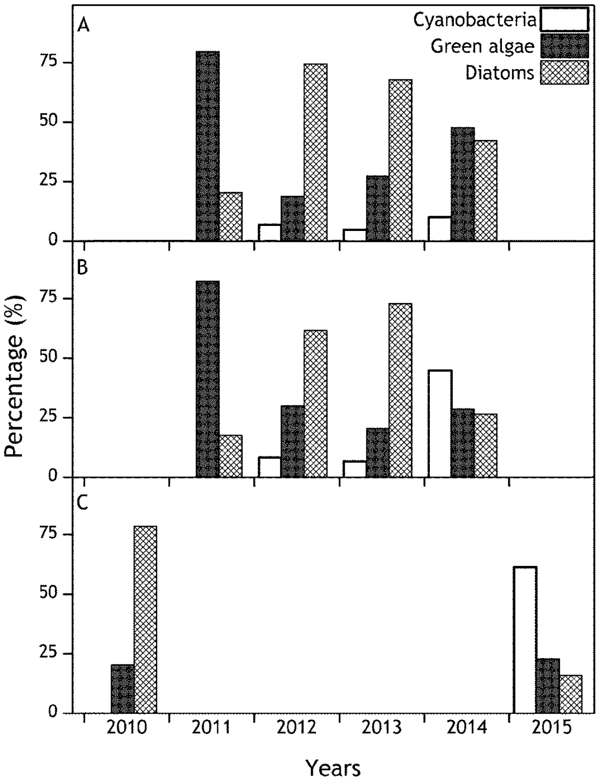

Fig. S4: Vertical oxygen profiles of both basins at Gollinsee on three dates (A: 17/07; B: 25/09; C: 25/11) in summer and autumn of 2013.

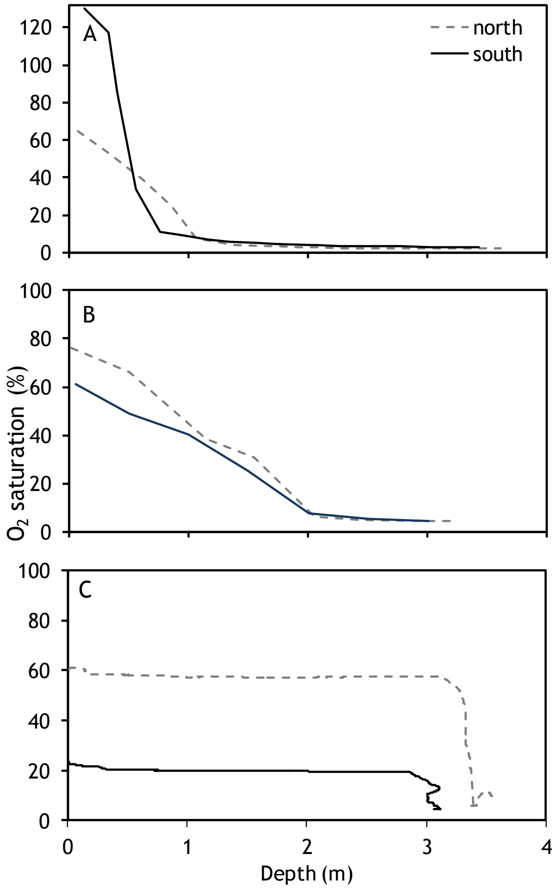


Fig. S5: The light extinction coefficient ($K_D$ in $m^{-1}$) in Gollinsee as a function of different DOC concentrations (black circles, mg $L^{-1}$). The dotted line represents $K_D$ used in the model of Kelly et al. (2018).

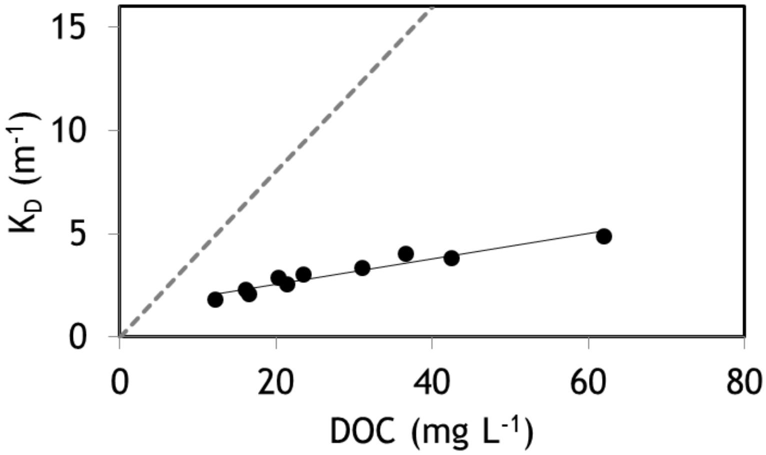
